# Do demographic processes change at extremely low population size in western monarch butterflies?

**DOI:** 10.1101/2021.10.22.465529

**Authors:** Collin B. Edwards, Cheryl B. Schultz, Elizabeth Crone

## Abstract

Allee effects – the breakdown of biological processes at low population densities – are ecologically important because they can potentially drag already struggling populations to extinction. However, identifying and documenting Allee effects is challenging, especially for natural populations, because it is difficult to know when populations have dropped to critically low densities, and to observe them both above and below this threshold. Here, we compared demographic processes in the western monarch butterfly, *Danaus plexippus,* before and after the population had fallen below the size at which Allee effects were hypothesized to take hold. Comparisons drew on data we collected after a dramatic population crash in 2018, previously published data from other researchers, and community science data. We found no evidence for Allee effects in winter survival, the fraction of females mated in early spring, or eggs laid per day. We did identify a 43% decline in the distance of seasonal range expansion, which could reflect Allee effects in terms of summer population growth rates or density-dependent movement behavior. In addition, overwinter survival of western monarch butterflies has substantially declined since first estimated in 1975 and may be contributing to the long-term population decline. The lack of evidence for Allee effects and the recent rebound in population size provide a more hopeful view for monarch conservation in the future but do not supersede the documented density-independent population decline across the last several decades.

## Introduction

Positive density dependence at low population size – known as Allee effects – can lead to rapid decline and extinction in populations that had previously exhibited only gradual declines or stationary but stochastic fluctuations in population size (Stephens et al. 1999, Courchamp et al. 1999). Allee effects can also limit range expansion and biological invasion, as individuals at range edges and rare long-distance dispersers can experience low local population densities (Taylor and Hastings 2005). When a population falls to sufficiently low density (below it’s “Allee threshold”), vital biological processes like mate-finding, group dynamics, and predator satiation can break down (Liermann and Hilborn 2001, Tobin et al. 2007), further reducing population growth rate. The most direct evidence for Allee effects thus comes from signals of density dependence in population growth rates at low densities.

For natural populations, many past studies have tested for positive density dependence at low population sizes by analyzing time series of a single population or comparing growth rates across populations. However, direct evidence of reduced growth rates is hard to obtain in practice (Kramer et al. 2009, but see Budroni et al. 2014, Shuai et al. 2020, Li et al. 2022). To identify Allee effects from time series data, ecologists need sufficient statistical resolution to distinguish Allee effects from year-to-year variability and other determinants of growth rate. Decomposing population growth rates into individual drivers is a considerable task under the best circumstances. When looking for Allee effects, ecologists face two additional challenges. First, Allee effects operate over a range of low population densities in which logistical challenges, measurement error, and sometimes conservation concerns can make it difficult to quantify population change. Second, under the conditions when Allee effects should occur (small populations), the stochasticity associated with small populations and the Allee effect itself (if present) can accelerate population decline (Liermann and Hilborn 2001). In combination, ecologists have limited ability to identify a signal of Allee effects, and frequently have limited time to do so before the population is extirpated.

An alternative approach to identifying Allee effects is to instead identify positive density dependence in individual biological processes: “component Allee effects” (Stephens et al. 1999). These component Allee effects are distinct from positive density dependence in overall population growth rates (hereafter “demographic Allee effects,” following Stephens et al. 1999), and component Allee effects are sometimes compensated for by negative density dependence in other processes. Component Allee effects are often more experimentally tractable to evaluate than demographic Allee effects (Kramer et al. 2009). Studies of component Allee effects measure how some essential process (often fecundity, foraging, or survival) varies between groups of individuals that, either by chance or through manipulation, experience different conspecific densities. These can include studies in artificial settings (Elliott et al. 2017, Ventura et al. 2017, Stringer et al. 2020) or short-term observational studies (McInnes et al. 2017, Ventura et al. 2017, Bayer et al. 2018, Contarini et al. 2009). One limitation of many of these studies is that it is challenging to know whether artificial settings accurately mimic low population sizes in the field, and whether spatial variation in abundance reflects factors other than population sizes. While rare, a few studies test for component Allee effects are based on long-term data, including estimating density dependence in reproduction rates of the crested Ibis (Li et al. 2022) and the dwarf-striped hamster (Shuai et al. 2020) and nest survival for Oystercatchers (Frauendorf et al. 2022).

In recent years, the migratory population of western monarch butterfly (*Danaus plexippus*) has undergone dramatic fluctuations in abundance, creating an opportunity to test for Allee effects. As part of a population viability analysis (Schultz et al. 2017), the authors used expert opinion to estimate that Allee effects would lead to population declines when the population dropped below 30,000 overwintering individuals. After declining at a rate of about 6% / year from the 1980’s (millions of butterflies) through mid-2010’s (∼200-300 thousand butterflies; Schultz et al. 2017), the population experienced a precipitous 85% decline in 2018-2019 to fewer than 30,000 butterflies (Pelton et al. 2019). The population had ∼20,000 butterflies in 2019-2020, and fewer than 2,000 butterflies counted in 2020-2021. It then experienced dramatic growth to more than 200,000 butterflies in 2020-2021 and modest growth in 2021-2022, ending 2022 with an overwintering population of approximately a third of a million (Xerces Society Western Monarch Count 2022). Despite the more hopeful current situation, between spring 2018 and fall 2021, there were very real fears that the migratory population of western monarch butterflies might not persist. From an academic perspective, this situation was a rare opportunity to evaluate what happens to a natural population when it drops below a hypothesized Allee threshold. These circumstances are rare, and studies taking this approach are similarly rare (but see Berec et al. 2007). Although it is now known that the migratory population of western monarch butterfly was not destined for extinction after the crash in 2018-2020 as it escaped from low population size, it is informative to find out how demographic processes changed during the low population years.

In this study, we evaluate changes in five processes and parameters that relate broadly to vital rates (survival, reproduction, and movement): (1) winter survival, (2-5) body size, wing size, matedness, and fecundity of butterflies at the end of winter dormancy, and (6-7) population range size and rate of expansion during summer breeding. Western monarch butterflies are characterized by seasonal multi-generational range expansion, in which the population begins the spring in a series of overwintering sites along the coast of California, expands into summer breeding grounds throughout western North America across several generations, then returns to the overwintering sites again in the fall, where they spend the winter in reproductive diapause (Dingle et al. 2005). Our focus was primarily on overwintering processes (metrics 1-5 in the list above) because of extensive past research on overwintering monarch butterflies in California (see Table 1), with an emphasis on the benefits of aggregation during this time period. For example, Anderson and Brower (1993) hypothesized density dependence in thermoregulation during the winter, suggesting Allee effects on winter survival or body size at the end of winter. Wells et al. (1990) and Wells et al. (1998) hypothesized density dependence in mate-finding for monarchs during the period of mating before aggregations break up in spring, suggesting Allee effects on matedness or fecundity.

**Table 1.** Sources of data for comparing demographic processes and patterns. Subscripts indicate geographic locations of data coverage.

|  |  |  |  |
| --- | --- | --- | --- |
| Process / pattern | Data sources for different eras |  |  |
|  | Historic | Declining | Post-crash |
| 1. Winter survival | Tuskes & Brower 1978<sup>1,2</sup> | Western Monarch Count<sup>9</sup> | Western Monarch Count<sup>9</sup> |
| 2. End-of winter body mass | Chaplin & Wells 1982<sup>5</sup>; Leong et al. 1995<sup>4</sup> |  | 2019 data (this study)<sup>4</sup> |
| 3. End-of-winter wing size | Beall 1946<sup>6,7,8</sup>; Tuskes & Brower 1978<sup>1,2,3</sup>; Leong et al. 1995<sup>4</sup> |  | 2019 data (this study)<sup>4</sup> |
| 4. End-of-winter matedness | Leong et al. 1995<sup>4</sup> | Leong et al. 2012<sup>4</sup> | 2019 data (this study)<sup>4</sup> |
| 5. End-of-winter fecundity |  | Leong et al. 2012<sup>4</sup> | 2019 data (this study)<sup>4</sup> |
| 6. Summer range size |  | Western Monarch Milkweed Mapper<sup>9</sup> | Western Monarch Milkweed Mapper<sup>9</sup> |
| 7. Rate of summer range expansion |  | Western Monarch Milkweed Mapper<sup>9</sup> | Western Monarch Milkweed Mapper<sup>9</sup> |
| <sup>1</sup>Muir Beach (37.86°N, 122.41°W); <sup>2</sup>Natural Bridges (37.95°N, 122.06°W); <sup>3</sup>Santa Barbara (34.41°N, 119.87°W); <sup>4</sup>Pismo Beach (35.07°N, 120.37°W); <sup>5</sup>Rancho dos Pueblos (Goleta) (34.44°N, 119.96°W); <sup>6</sup>Ventura (34.28°N, 119.29°W); <sup>7</sup>Pacific Grove (36.62°N, 121.92°W); <sup>8</sup>also includes some samples from Bolinas, El Cerrito, San Francisco, San Jose, Owens Valley and San Diego; <sup>9</sup>range-wide data for western monarch butterflies |  |  |  |

During summer breeding, past research on monarch butterflies has tended to emphasize the costs of crowding, i.e., negative density dependence (Fisher & Bradbury 2022, Flockhart et al. 2012, Lindsey et al. 2009, Marini and Zalucki 2017, Yakubu et al. 2004), as opposed to positive density dependence and possible Allee effects. However, it is reasonable to expect that density-dependent mate-finding could be at least as problematic during summer breeding, when populations can be distributed sparsely over large geographic areas, as during mating at the end of overwintering aggregations. Although Allee effects have not specifically been proposed for monarch butterflies during summer range expansion, small populations at geographic range margins are a classic example of potential Allee effects in theoretical models, and it is well known that small population sizes can slow or prevent population expansion (e.g., Taylor and Hastings 2005, Lewis and Kareiva 1993, Hastings 1996, Morel-Journel et al. 2019). To our knowledge, no prior studies have measured western monarch matedness or female fecundity during summer breeding before the population crash. Therefore, we used community science observations to analyze breeding season range expansion – the rate at which populations expand into summer breeding grounds – as a metric of demographic processes during this time period (Table 1). Rates of geographic range expansion depend on both population growth rates and movement (for a textbook-level introduction see, e.g., Kot 2001). Therefore, changes in monarch seasonal range expansion after the population crash could reflect changes in survival, reproduction or movement rates (cf. Crone and Schultz 2022).

For each of the seven processes and patterns, we expect that, if component Allee effects are present (and we correctly estimated the Allee threshold), these measures would be lower during and after the 2018 crash than during prior time periods. However, many factors other than population size are also likely to vary through time, especially when comparing metrics from the mid-20^th^ century to the post-crash period. Therefore we divided data into three broad time periods: “historical,” before 2000, when there were millions of monarchs; “declining,” from 2000-2017 when monarchs were in the hundreds of thousands and slowly declining; and “post-crash,” from 2018-2020 when winter counts were below the hypothesized Allee threshold. We note that our eras are based on population sizes, with labels that are generally descriptive rather than prescriptive. For example, we know declines occurred in the late historical era, and the population in the declining era sometimes increased in size in individual years. In our analyses, differences between the post-crash era and the declining era indicate the presence of Allee effects. Differences between the historic era and post-crash or declining eras indicate possible drivers of long-term declines.

## Methods

### Study system

The monarch butterfly (*Danaus pelxippus*) is a broadly distributed species, originating from the Americas, that has colonized many parts of the world, including populations in Europe, Africa, Australia, and numerous Pacific and Caribbean islands (Agrawal 2017). In North America, there are two migratory populations, a larger eastern monarch population that overwinters in Mexico and breeds in North America east of the Rocky Mountains and a smaller western population that overwinters on the California coast (the state of California in the USA and northern Baja California in Mexico) and breeds in North America west of the Rocky Mountains. These two migratory populations are not strongly genetically differentiated, but they are demographically distinct, in the sense that they fluctuate independently (Freedman et al 2021). Although both populations have experienced declines in recent years, the western population is both smaller and is declining at a faster rate, so is at greater risk of extirpation (Semmens et al. 2016, Schultz et al. 2017).

The western monarch butterfly has a multi-generational life cycle: overwintering individuals emerge from reproductive diapause and travel from the coast of California to lay eggs before they die; many of their offspring also fly substantial distances before laying eggs, and across three to four generations each summer the butterflies spread throughout the western United States and into southern Canada (Dingle et al. 2005). The final generation flies back to overwintering sites on the coast of California in the fall, completing the cycle. Causes of long-term population declines in western monarch butterflies include urban development and degradation of coastal overwintering sites (Pelton et al. 2019), as well as climate change (Stevens and Frey 2010, Espeset et al. 2016, Crone et al. 2019), increased pesticide use (Crone et al. 2019, Halsch et al. 2020), and declines in abundance of its milkweed (*Asclepias* spp.) hostplants (Pelton et al. 2019, Crone and Schultz 2022).

Our inference about patterns of abundance in the “eras” defined in our study comes from two sources. The migratory population of western monarchs is monitored annually at wintering grounds using a systematic community science survey, the Western Monarch Count, which includes “Thanksgiving Counts” (in late November and early December, 1997-present) and the “New Year’s Counts” (in late December and early January, 2017-present) (Xerces Society Western Monarch Count 2022). Schultz et al. (2017) combined these data with similar counts at wintering grounds to extend this time series to include population estimates from the 1980’s through 2016. These data provide the general population estimates that define our demographic eras. In addition, Schultz et al. (2021) monitored monarch butterfly breeding abundance and phenology at 12 sites in five western states (Washington, Oregon, California, Idaho and Nevada) from 2017-2019, and 2023-ongoing. These data showed that the population crash in the 2018 Thanksgiving count occurred in early spring of 2018 (Pelton et al. 2019). Therefore, we include the summer of 2018 in the “post-crash” era during analyses of breeding season dynamics.

### Changes in processes and parameters at wintering grounds

#### Winter: pre-crash studies

To estimate historic era survival, we used analyses presented by Pelton et al. (2019). This previous analysis used mark-recapture data presented in Tuskes and Brower (1978) to estimate 6-week survival for western monarchs in 1975-1976 near Santa Cruz and at Muir Beach (6 weeks is approximately the time between the Thanksgiving and New Years Counts of the Xerces society). For this paper, we combined the survival estimates for the two locations. In doing so, we corrected one typo in Pelton et al. (2019): the “29% drop” reported for Santa Cruz should instead say “19% drop”.

For other winter processes and patterns, we identified studies in the historical and declining eras that directly or indirectly report data relating to winter survival, early spring fecundity, early spring matedness, early spring body size, or wing size (Table 1). Although body size and wing size are not direct demographic measures, they were included because they often relate to survival, fecundity and movement ability (for which we had limited or no data). Where possible, we extracted the raw data from published tables and carried out our own analyses as described below. Where this was not possible, we extracted means and standard deviations from published figures using WebPlotDigitizer (Rohatgi 2021). When more than one data set was available for the same era, we combined the estimates, propagating uncertainty using the delta method (Williams et al. 2002), implemented in the R package msm (Jackson 2011). The delta method is a flexible mathematical tool for estimating the variance of a derived parameter estimate (e.g., the average of two or more parameter estimates from different studies, each with its own variance).

To estimate historical body size, we digitized means and standard errors of body mass data for female western monarchs from Chaplin and Wells (1982) (data from Pismo Beach in February 1972) and Leong et al. (1995) (data from Pismo Beach in February in years 1990-1992). To estimate historical wing size, we digitized data from Beall (1946), Tuskes and Brower (1978), and Leong et al. (1995), who reported estimates for wing length of female western monarchs caught primarily from Ventura and Pacific Grove in Februaries 1938-1939 and 1942, from Muir Beach, Natural Bridges and Santa Barbara in February 1976, and Pismo Beach in Februaries 1990-1992 respectively. Leong et al. (1995) caught 142 females in February of 1990, 1991, and 1992 at Pismo Beach and identified mating status using dissections and spermatophore identification; Leong et al. (2012) did the same with 19 females caught in February 2010. We fit these two data sets separately using an intercept-only binomial regression model to estimate the proportion of matedness in the historical and declining eras. Leong et al. (2012) reported estimated means and standard deviations for the lifespan (N=19) and lifetime eggs per female (N=10) for monarchs in February 2010. We calculated mean eggs per day as mean lifetime eggs divided by mean lifespan, using the delta method to propagate error.

#### Winter: post-crash data & analyses

We used changes in the number of monarch butterflies at overwintering sites to estimate winter survival. Starting in January 2017 (winter 2016-17), the Xerces Society carried out surveys in early January (“New Years Count” data) to complement their Thanksgiving Count surveys (Xerces Society Western Monarch Count 2022). We used binomial regression to estimate winter survival for this period as the ratio of the summed New Year’s Count (events) to the Thanksgiving Count (trials) for each year from winter 2016-2017 through winter 2020-2021. We fit these five years of data in a single binomial regression model, with a fixed effect of era (“declining” for the first two winters and “post-crash” for the most recent three) and a random effect of year.

We estimated other winter processes and patterns in a post-crash field study. (Due to low population sizes, we were not able to obtain permits to repeat Tuskes’ and Browers’ mark-recapture study to also measure survival directly.) In February 2019, we collected 35 adult female monarchs near the overwintering site at Pismo Beach State Park. We measured wing length and body size, and estimated means and standard errors using intercept-only regression models. We also used the monarch spermatophore palpation method to evaluate whether butterflies collected had been mated (Van Hook 1999). We treated females with a palpation score of 1 (“rice grain or large pinhead”) or higher as having successfully mated, and females with scores of 0 as unmated. Two individuals had an uncertain score and were excluded from further analysis. We used an intercept-only binomial regression model to estimate the proportion of mated individuals. Finally, to estimate daily fecundity, we kept adult female monarchs from Pismo Beach in tents in the greenhouse for two days with access to narrowleaf milkweed plants (*Asclepias fascicularis*), nectar feeders (sponges soaked in Gatorade) and nectar plants (*Lavandula*, *Iberis*, and *Chrysanthemum* sp.) sourced from local garden stores. At the end of two days, we counted all eggs laid by each female during this time and released butterflies back into the site from which they were collected. We analyzed these oviposition data using 0-inflated negative binomial models (using the pscl package in R) (Zeileis et al. 2008, Jackman 2020). 0-inflated models are a statistical technique that separates 0’s (in this case, butterflies that were not reproductively active) from a process that might include 0’s and higher values (in this case, eggs laid per day for reproductively active butterflies). The model produces two parameters: the proportion of 0’s (hereafter, the proportion of non-reproductive butterflies) and the mean of the non-zero distribution (here, the average count of eggs per reproductive butterfly). Unlike Leong et al. (2012), we did not carry out manipulations to break our captured monarchs from reproductive diapause. Therefore, 0’s from the 0-inflated negative binomial models likely represent butterflies in reproductive diapause, but could also represent unmated butterflies, and we do not interpret this parameter further.

#### Winter: comparing eras

To compare differences between eras, we carried out parametric bootstrapping. We generated 1000 replicates of estimated values, generally by drawing randomly from the distribution associated with the combined historical data or from the distribution of the appropriate coefficient estimate from fitted regression models using the bootMer() function of the R package lme4 (Bates et al. 2015). For matedness in the declining era, 100% of individuals were mated, which produces unreasonable estimated distributions using logistic regression. In this case, we sampled directly from the likelihood profile using rejection sampling.

### Changes in summer population expansion

#### Summer: estimating range expansion

We obtained observations of monarchs in California, Oregon, Washington, Nevada, and British Columbia from the late 1800s through January 1, 2023, from iNaturalist (GBIF.org 2020) and the Western Monarch Milkweed Mapper (Western Monarch and Milkweed Occurrence Database 2018); these observations included latitude, longitude, and day of year of observation. Most analyses in this paper are based on data only through December 2020. We carried out additional analyses of 2015-2022 to identify more precisely define the fingerprint of the decline and the 2021 rebound in abundance (GBIF.org 2023).

Because there are resident, non-migratory populations of monarch butterflies in Southern California and southern Nevada, we removed all observations south of the line drawn by connecting the locations (−120.63 latitude, 34.75 longitude) and (−114.02 latitude, 38.49 longitude) (Wells and Wells 1992) (Fig. 2A). We also excluded observations less than 30 kilometers from the coast, as some fraction of individuals have recently begun to remain on the coast or in coastal cities (notably, San Francisco and its environs) year-round (Crone and Schultz 2021). The specific distance cutoff used for exclusion did not affect our conclusions (see “*Summer: sensitivity analysis*” below). We removed duplicate observations (defined as having the same date and latitude and longitude each within 1/1000 degree of each other), and removed obvious outliers based on expert opinion (36 outliers total). For each of the remaining 4489 observations, we calculated the great circle distance (“as the crow flies”) to the nearest overwintering sites, using the 185 overwintering sites of Schultz et al. (2017). For this analysis, we used summer 2018 as the start of the post-crash era because the low abundance started in spring 2018 (Pelton et al. 2019). Analyses of individual years from 2015 to 2022 used the same methods and data sources.

Plots of observations based on day of year and distance from the nearest overwintering sites showed a roughly symmetrical, hump-shaped distribution that could be approximated by a Gaussian curve (e.g., Fig. 2B). Edwards and Crone (2021) show how Gaussian curves can be fit with linear models; we combine this approach with quantile regression to estimate the outer boundary of monarch seasonal range expansion using the quantreg package (Cade and Noon 2003, Koenker 2021). We used the 0.9 quantile to represent the outer boundary of the monarch range; this defines the boundary between the 90% of monarchs nearest the overwintering sites and the most distant 10% on any given day of the year. We chose this method because it captures the expansion of monarchs into their summer breeding range while still being robust to outliers. The specific quantile chosen did not affect our conclusions (see “*Summer: sensitivity analysis*” below). For comparison to analyses of winter data, we aggregated our data into the same historical, declining, and post-crash eras before fitting our models. From fitted curves of the 0.9 quantile for each era, we estimated the maximum distance from overwintering sites as a metric for the summer breeding range size. As metric of the rate of range expansion, we estimated when the 90% quantile first reached 100 kilometers from the nearest overwintering site. One hundred kilometers is approximately the (greater circle) distance from the nearest overwintering site to Sacramento, California’s state capital, and was well within the maximum distance traveled in any era. For both metrics (maximum distance and days to 100 km), we estimated confidence intervals using bootstrapping as implemented in the msm package (Jackson 2011). All calculations were based on day of year (0-366); for readability, we present our results as month and day for a non-leap-year.

As we had sufficient data to fit separate models to each year for 2015-2022, we repeated our calculations on a yearly basis for this period. Our yearly fits allowed us to examine seasonal range expansion before the 2018-2019 crash, after the crash, and then after the 2020 rebound in abundance.

#### Summer: sensitivity analysis

To confirm that the results of our range expansion analysis did not depend on specific model-fitting decisions, we repeated the main analysis with each of the following changes: defining the monarch range by the 0.8, 0.7, 0.6, and 0.5 quantiles (instead of the 0.9 quantile used above), or changing the threshold distance from the coast below which we excluded points to 0.1 km, and then 50 km (instead of 30 km used above). Additionally, we observed apparent streaks in the raw data when plotting distance by day of year (Fig. 2B, Fig. S4). The highly overrepresented distances may reflect repeated sampling of populated or popular locations. To ensure these streaks were not driving observed patterns, we generated simulated data sets by randomly subsampling our data such that we never had more than 1 data point in each 1km distance from the overwintering sites for each era. This sampling procedure generated simulated data sets with 1276 observations instead of 4492. We repeated this process 5000 times and carried out our previous analysis on each of these simulated datasets.

### Software and key packages

All analyses and figure generation were carried out in the programming language R version 4.2.1 (R Core Team 2022). We used the following key packages: spdata (Bivand et al. 2023), geosphere (Hijmans 2022), maps (Becker et al. 2022), mapdata (Becker and Wilks 2022b), and ggmap (Kahle and Wickham 2013) for our spatial analyses and plots; lme4 (Bates et al. 2015) for regression and mixed models, pscl for zero-inflated regression (Jackman 2020), quantreg for quantile regression (Koenker 2022); msm (Jackson 2011) and boot (Canty and Ripley 2022) for bootstrapping; tidyverse suite (Wickham et al. 2019) for data manipulation; ggplot2 (Wickham 2016), scales (Wickham and Seidel), gridExtra (Auguie 2017), and patchwork (Pederson 2022) for figure-making. For supplementary animations, we used the r “animation” package (Xie et al. 2021) and the software ImageMagick (ImageMagick Studio LLC. 2023).

## Results

None of the demographic processes and patterns measured at the wintering grounds showed evidence of component Allee effects. Winter survival was significantly higher in the post-crash era than in the declining era (higher in 97.2% of bootstraps), indicating no Allee effect on this process (Fig 1A). Winter survival was significantly higher in the historical era (75.4%, 95% CI = 71.2% -- 79.0%) than in the declining (53.7%, 95% CI =48.4% -- 58.9%) or post-crash eras (60.4%, 95% CI = 56.2% -- 64.6%) (higher in 100% of bootstraps in both cases). Body mass was slightly higher in the post-crash era (0.48 grams, 95% CI = 0.48 – 0.508) than in the historical era (0.47 grams, 95% CI = 0.461 – 0.478), but this difference was not statistically significant (higher in 83.5% of bootstraps), and was in the opposite direction of an Allee effect (Fig 1B). Wing length was greater in the post-crash (52.4 mm, 95% CI = 51.4--53.4) than historical era (51.2 mm, 95% CI = 50.8 – 51.6) (higher in 98.1% of bootstraps) (Fig. 1C). The fraction of females mated was highest in the declining era (100%, 95% CI = 90.3% -- 100%), followed by the post-crash era (88%, 95% CI = 74% -- 96%) and then the historical era (70%, 95% CI = 63% -- 78%) (Fig. 1D). The fraction of females mated was significantly higher in the post-crash and declining eras than the historical era (higher in 97.8% and 100% of bootstraps, respectively), while the difference between declining and post-crash eras was not significant, indicating there was no support for a component Allee effect (declining era had higher matedness in 87.7% of bootstraps). Egg production was higher in the declining era (12.8 eggs per day, 95% CI = 3.3 – 22.4) than in the post-crash era (5.5 eggs per day, 95% CI = 2.7 – 11.2), but this difference was not statistically significant (declining era was higher in 90.2% of bootstrapped replicates) (Fig. 1E).

**Figure. 1.**
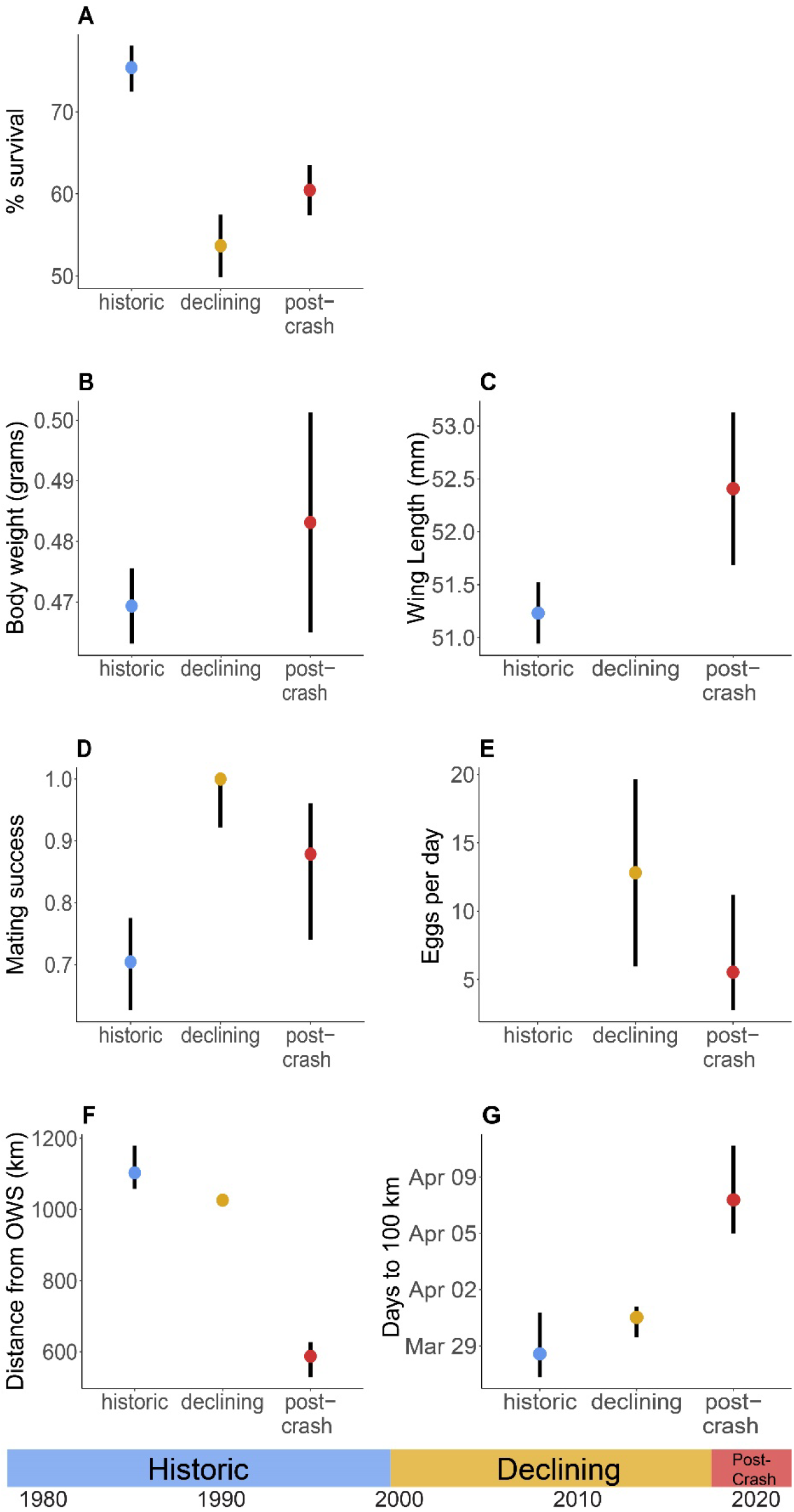
Comparison across eras for (**A**) winter survival (from Thanksgiving to New Year’s Day), (**B**) body mass, (**C**) wing length, (**D**) proportion of females mated, (**E**) fecundity, (**F**) summer breeding season range as measured by maximum distance from overwintering sites, and (**G**) the day the 0.9 quantile of seasonal range expansion reaches 100 km from overwintering sites (OWS). Points correspond to estimated values before 2000 (“historic”), from 2001-2017 (“declining”), and from 2018-2020 (“post-crash”) when the population was below the hypothesized Allee effect threshold. Bars represent 84% confidence intervals (non-overlapping bars correspond to significant 2-tailed T tests; Payton et al. 2003).

There was strong evidence for positive density dependence in monarch summer breeding range size and the rate of range expansion. The historic and declining eras had similar summer breeding ranges, with the 0.9 quantile of the population expanding to up to 1103 km (95% bootstrapped CI = 1037 –1202) and 1001.98 km (95% bootstrapped CI = 1002 – 1051) from overwintering sites, respectively (Fig. 1F, Fig. 2B, Movie S1). This 9% decline in distance from the historic to the declining eras was statistically significant (historic era distances were greater than declining era distances in 98.5% of bootstrapped replicates). In the post-crash era, the maximum distance traveled fell by a further 43% to 587 kilometers, suggesting density-dependent range expansion (95% bootstrapped CI = 518 – 657 km; less than historical and declining eras in 100% of bootstraps). Models fit separately to each year of 2015-2022 show a clear relationship between breeding range size and population size, with the distance of range expansion falling in 2018 at the time of the population crash, then increasing somewhat during and after the 2020 bounce (Fig. 2C).

**Figure. 2.**
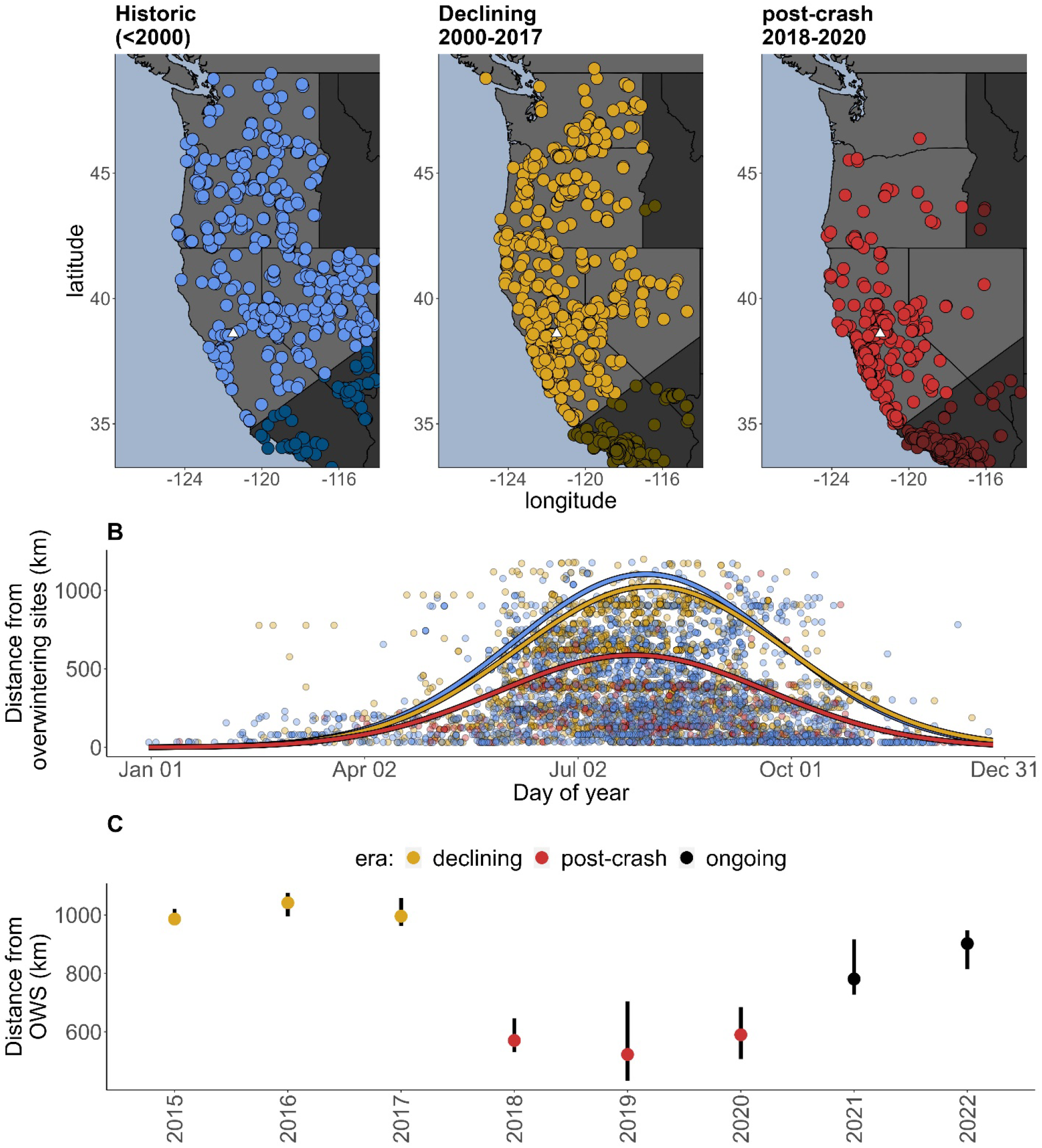
(**A**) Fewer monarchs were observed in OR and WA in the post-crash era (shaded CA and NV regions were excluded to avoid resident populations). White triangles show Sacramento, CA, a representative location 100 km from the overwintering sites. (**B**) Fitted curves of the 0.9 quantile of the distance from observations to the nearest overwintering site show substantial reduction and an earlier peak in the post-crash era, and are robust to differences in abundance. (**C**) High numbers of monarch observations allow us to fit the past eight years individually, calculated as in Fig. 1F. The dramatic decline in maximum distance from overwintering sites (OWS) exactly corresponds to the crash in early 2018; the bounce in 2020-2021 corresponds to a partial recovery of the maximum distance.

The rate of range expansion showed similar patterns (Fig. 1G): Monarch populations reached 100 km from the coast at a slightly earlier date in the historic (March 28, 95% bootstrapped CI: March 26-April 1) and declining (March 31, 95% CI: March 29 – April 1) eras (declining era later than historic in 87.4% of bootstraps). In the post-crash era, the date at which they reached 100 km from the coast was more than a week later than in the historic era (April 7, 95% CI: April 4-12; later than historical and declining eras in 100% and 99.9% of bootstraps).

When repeating our analyses using different quantiles and different distances for excluding coastal sightings, individual estimates changed, but the same qualitative patterns remained (Supplemental Fig. S2, S3). Similarly, randomly subsampling our data to remove streakiness changed specific estimates but produced qualitatively similar results to our analysis of the complete data set (Supplemental Fig. S4).

## Discussion

We found no evidence for component Allee effects in demographic processes or patterns for monarch butterflies during winter. Matedness and fecundity were lower in the post-crash era than in the declining era, but these differences were not statistically significant; overwintering survival was significantly higher in the post-crash era than in the declining era.

We expected to find evidence for mate limitation, as this is one of the best-documented component Allee effects in other species (Kramer et al. 2009, Gascoigne et al. 2009) and has been specifically identified as a potential issue for overwintering monarch butterflies (Wells et al. 1990, Wells and Wells 1992). Similarly, overwintering aggregation has long been proposed as a method for temperature regulation to improve survival (Allee 1927), and has also been hypothesized as a benefit of monarch aggregation during overwintering (Anderson and Brower (1993), although Chaplin and Wells (1982) found that overwintering monarchs were generally at air temperature regardless of their level of aggregation, and Wells and Wells (1992) outline further indirect evidence that winter aggregations do not mitigate temperature-related mortality.

Certainly, one possibility is that our historical data were too sparsely distributed in space and time (see Table 1) to provide a robust comparison with our estimates from 2019. An ideal but unrealistic experimental design would be to measure vital rates at multiple sites before and after the population crash; this study design is unrealistic because we could not have predicted this population crash and also because there were too few butterflies at most sites to conduct research after the crash. A similar concern is that we did not know the correct threshold population size for demographic Allee effects; our study design was based on a published Allee threshold (Schultz et al. 2017), but the exact threshold value was largely a guess based on expert opinion. Furthermore, in other systems, threshold population sizes for demographic Allee effects are known to vary spatially and temporally (Tobin et al. 2007, Walter et al. 2020), so our estimated Allee effect threshold may have been correct on average, but fortunately in this instance the population avoided demographic Allee effects. Nonetheless, in spite of these caveats, our study was a rare opportunity to repeat past demographic studies after a sudden 85% drop in population size.

An alternative “null” explanation for lack of component Allee effects during overwintering is that the individual site surveyed did not reach low enough densities, even in this unprecedented population crash. Our survey and both past studies of matedness (Leong et al. (1995) and Leong et al. (2012)) were all carried out at Pismo Beach, in part because it is a reliable overwintering site with high densities of monarchs. Pismo Beach experienced a substantial reduction in overwintering numbers between the surveys in the declining era, where the average count was ∼25,000 individuals (range: 12,284-38,438), and our post-crash experiments, when 3,092 individuals were reported in Fall 2018 (Xerces Society Western Monarch Count 2022). However, in the fall of 2018, Pismo Beach had the most reported individuals of all overwintering sites. In contrast, 47% of monitored sites with at least one monarch present had fewer than ten individuals reported, and a third of monarchs reported were in sites with less than 600 individuals. It is possible that we would have detected component Allee effects at some of the less densely populated sites, although there were so few monarchs there that experiments would not have been feasible. It is also possible that we might have detected component Allee effects in the winter of 2020-2021, when the overwintering population crashed again to an all-time low Thanksgiving Count of fewer than 2000 monarchs counted across all sites. However, at that point, there were essentially no overwintering sites with enough monarch butterflies to conduct rigorous experiments; in 2020 Pismo Beach had a Thanksgiving count of 199 butterflies, and only one site had a higher count (550 at Natural Bridges). Thus, our study is an unusual opportunity to test hypothesized component Allee effects, but also emphasizes the challenges of doing so (cf. Liermann and Hilborn 1997, 2001).

The most striking change between the declining and post-crash era was the nearly 50% reduction in the maximum distance of the seasonal range expansion. Demographic Allee effects slow the rate of geographic spread in mathematical and simulation models (Lewis and Kareiva 1993, Veit and Lewis 1996, Wang and Kot 2001, Morel-Journel et al. 2019), suggesting that the reduced rate of range expansion in western monarch butterflies could reflect lower survival or fecundity during breeding in the post-crash era. Past studies of monarch butterflies during the breeding season have shown either negative (Flockhart et al. 2012, Lindsey et al. 2009, Marini and Zalucki 2017) or no (Fisher & Bradbury 2022) density dependence. However, these studies were all done in the eastern monarch butterfly population, where monarch butterfly densities are typically much higher than in the West. Studies that experimentally manipulated densities also did not include very low densities, e.g., 1-10 larvae per 3.8L container (Lindsey et al. 2009) or 1-2 larvae per 4 milkweed (host plant) stems (Fisher & Bradbury 2022). In contrast, typical western monarch densities were one larva per ∼10-100 milkweed stems in 2017 (prior to the crash) and one larva per ∼1,000-10,000 stems in 2018 (after the crash) (Pelton et al. 2019). These past studies of density dependence also focused on competition at the life stages from oviposition through larval pupation. To our knowledge, no one has tested for density-dependent mate limitation in monarch butterflies during summer breeding, even though mate limitation was a leading reason why ecologists had proposed component Allee effects in overwintering aggregations (Wells et al. 1990, Wells and Wells 1992). Spongy moths, an invasive lepidopteran species, experience demographic Allee effects at their range edge due to mate-finding failures (Tobin et al. 2007, Contarini et al. 2009).

A second mechanism for reduced seasonal range expansion is density-dependent dispersal. Reduced dispersal at lower population sizes has been reported for several other butterfly species (Nowicki and Vrabec 2011, Bonelli et al. 2013, Kuussaari et al. 2016). In most cases, density dependent dispersal is framed as a way of avoiding crowded conditions. For the western monarch, an additional mechanism for positive density-dependent dispersal is a reduction in multiple matings of overwintering monarchs when the population is small (Wells and Wells 1992). Male monarch butterflies transfer a large nutrient-rich spermatophore during mating, and multiple matings can provide female butterflies with substantially increased energy, in turn increasing their dispersal capabilities (Wells et al. 1992, Wells and Wells 1992).

In contrast with the notion of reduced movement, monarch butterflies in our 2019 post-crash data set had larger wing sizes than samples from the historic era (Fig. 1C). We observed a 1.2 millimeter increase in size in the post-crash era, which is approximately equal to the difference observed between migratory and non-migratory individuals (Altizer and Davis 2010). Across monarch populations, larger wing sizes are associated with increased capability of movement: the eastern monarch butterflies, which migrate farther, have larger wings than western monarch butterflies or monarch butterflies from nonmigratory populations (Altizer and Davis 2010). Similarly, within the western monarch population, larger wings in overwintering individuals are associated with the isotopic fingerprints of more distant summer breeding grounds (Yang et al. 2016). Larger wing sizes could also be interpreted as a sign of more stressful conditions and higher mortality during Fall migration (Altizer et al. 2015), which could potentially translate to slower inland movement and/or lower fecundity during the first generation of monarch breeding in Spring. It is possible that small changes in the rate of range expansion during early generations could lead to mismatched timing of monarch butterflies and host plants, with knock-on consequences of reduced larval survival (cf. Yang and Cenzer 2020).

If reduced seasonal range expansion is a component Allee effect, it seems likely that other parts of the life cycle showed compensatory responses, so this effect did not lead to population declines. The western monarch population grew dramatically from Fall 2020 to Fall 2021, in spite of a continued reduction in the breeding range during the summer of 2021. Similarly, in Fall 2021 and 2022, the western monarch Thanksgiving count was similar to counts in the declining era, but the breeding range was significantly smaller in summer 2021 and summer 2022 than in the declining era, even though it was significantly larger than in the post-crash era. It is not uncommon for positive density dependence in one life cycle stage to be counterbalanced by demographic or behavioral changes at other life cycle stages (Levitan 1991, Shuai et al. 2020, Stephens et al. 1999). Some past studies of eastern monarch butterflies have suggested an opposite pattern of demographic compensation to our results for western monarchs: faster declines in winter than summer in the east (Inamine et al. 2016, Crossley et al. 2022), and faster recovery of winter population counts than breeding range size in our data. The difference in findings may arise in part because the summer counts used in Inamine et al. (2016) and Crossley et al. (2022) represent only a portion of the eastern monarch breeding range, and are likely biased toward higher-quality habitat (Pleasants et al. 2017). Additionally, Crossley et al (2022) compared trends in winter monarch counts for the eastern population to trends in summer monarch butterflies for all of North America, including western and non-migratory populations, so divergent trends might reflect different dynamics of these groups. Nonetheless, these studies all point to the general importance of considering Allee effects throughout a species’ life cycle, which includes understanding migratory species throughout their full annual life cycle.

In general, our results suggest that the western monarch butterfly population is remarkably resilient to fluctuations in abundance. This resilience may not be surprising, since insect populations often vary by orders of magnitude among years (Fox et al. 2019, Didham et al. 2020). However, this positive conclusion is moderated by at least three cautionary caveats: The first of these is that overwinter survival was much higher in the historic era than the declining and post-crash eras. This difference could be due to differences in methodology (mark recapture analyses in the historic era and inference from population counts in more recent years; see *Methods*). However, the western monarch population has declined substantially since the mid-20^th^ century and is about an order of magnitude smaller than in the 1980’s, even after the recent upswing in numbers. We emphasize the importance of conserving and restoring wintering groves, as well as summer breeding habitat (Pelton et al. 2019).

A second caveat is that, although the differences were not statistically significant, the estimated proportion of mated females and eggs laid per day fell in the post-crash era, consistent with component Allee effects. If these nonsignificant effects were real, females in the spring of the post-crash era would have laid less than half as many eggs on average as those in the declining era (% mated × eggs/day: declining era = 12.8; post-crash era = 4.9). This is within the range of other estimated component Allee effects on fecundity. In island foxes, the proportion of breeding females varied from approximately 60% at high-density sites to approximately 20% at low-density sites (Berec et al. 2007), the pregnancy rate of striped hamsters varied between approximately 35% and 10% from high to low density years (Shuai et al. 2020), and in two small reintroduced populations of Crested Ibis, the larger population averaged 65% larger clutch sizes and 58% higher breeding success (Li et al. 2022).

The third caveat is that, although the migratory population of the western monarch did not decline to extinction, we do not know how or why it increased from the Fall of 2020 to the Fall of 2021. The drivers of western monarch population dynamics are complex, interrelated, and ecologists are still seeking to understand them (Yang 2023). It might be that the western monarch population was suffering from severe component Allee effects in 2019 and 2020 but the detrimental consequences were counterbalanced by unusually good conditions or a “series of fortunate events” (see Greenberg 2022). There are many non-exclusive alternative explanations, including the potential that the population was rescued by migration from the east (there is at least some mixing between the populations) (Billings 2019). The lack of evidence for component Allee effects for the western monarch does not guarantee that the population will always recover from low abundance but does imply low population sizes are not a guarantee of extinction.

---

**Supplemental Figure S1:**
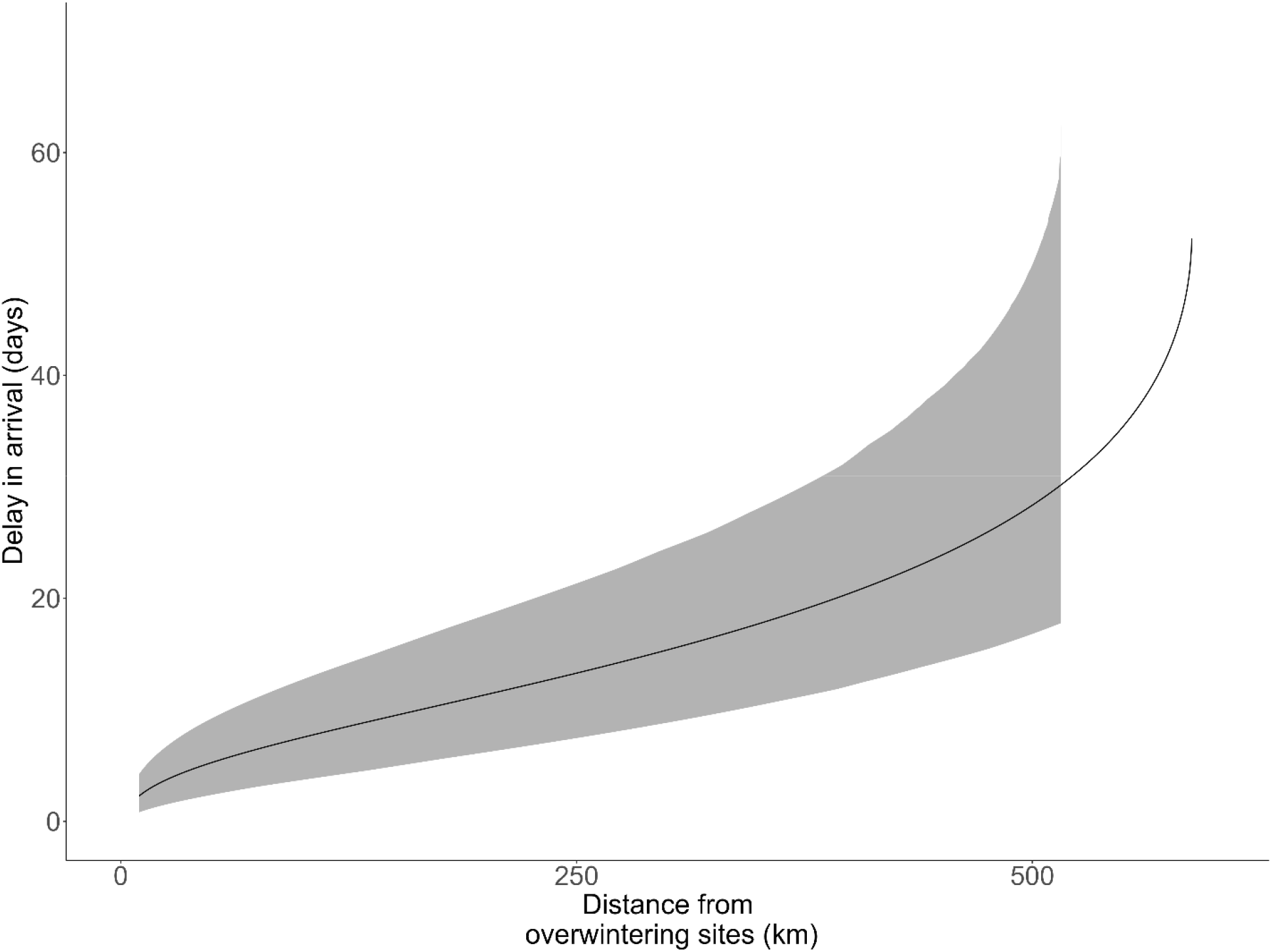
Difference in the days to 0.9 quantile between the declining and post-crash eras as a function of distance. Western monarch phenology was delayed in the post-crash era, and for regions sufficiently far from overwintering sites, the 0.9 quantile of seasonal range expansion arrived more than a month later in the post-crash era. Black line shows estimated distance, gray area shows bootstrapped 95% confidence envelope.

**Supplemental Figure S2:**
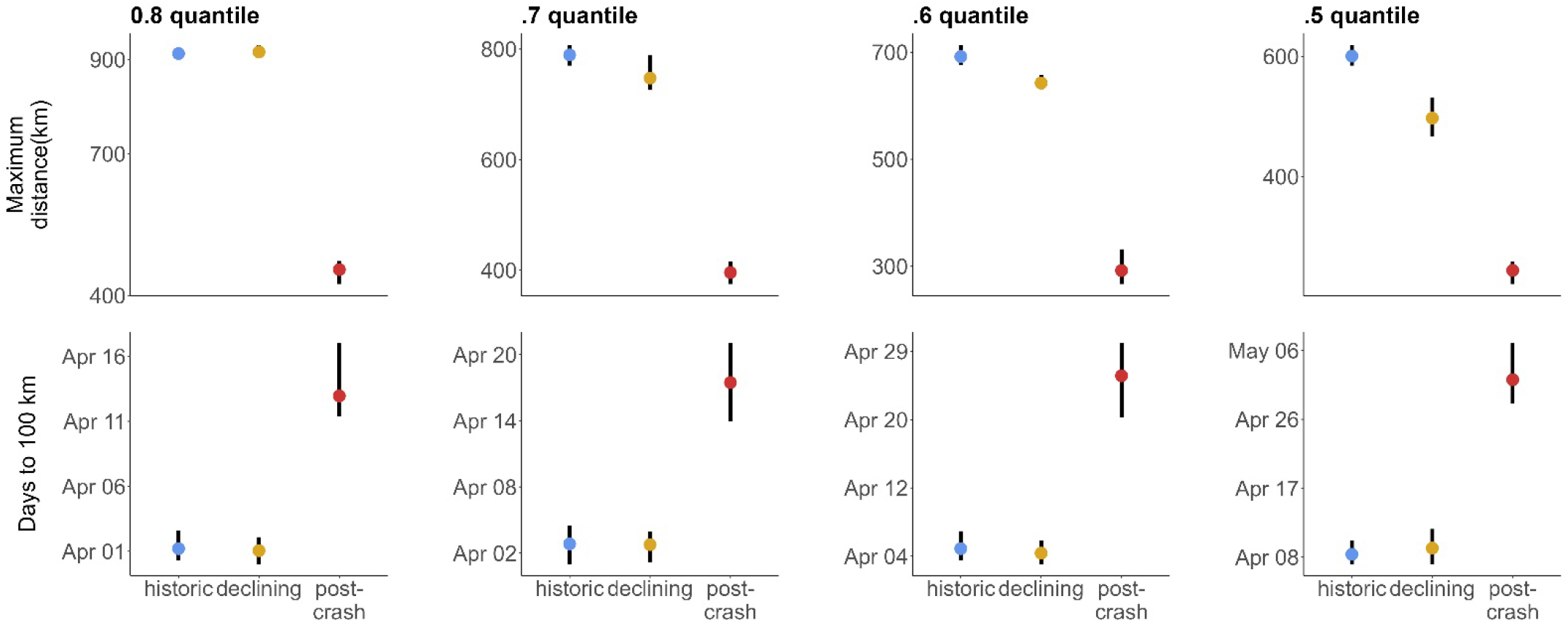
Qualitative patterns of seasonal range expansion are insensitive to changes in the quantile choice. We repeat our analyses and replicate Fig. 1F and 2C using 0.8 to 0.5 quantiles (as compared to 0.9 in the main text).

**Supplemental Figure S3:**
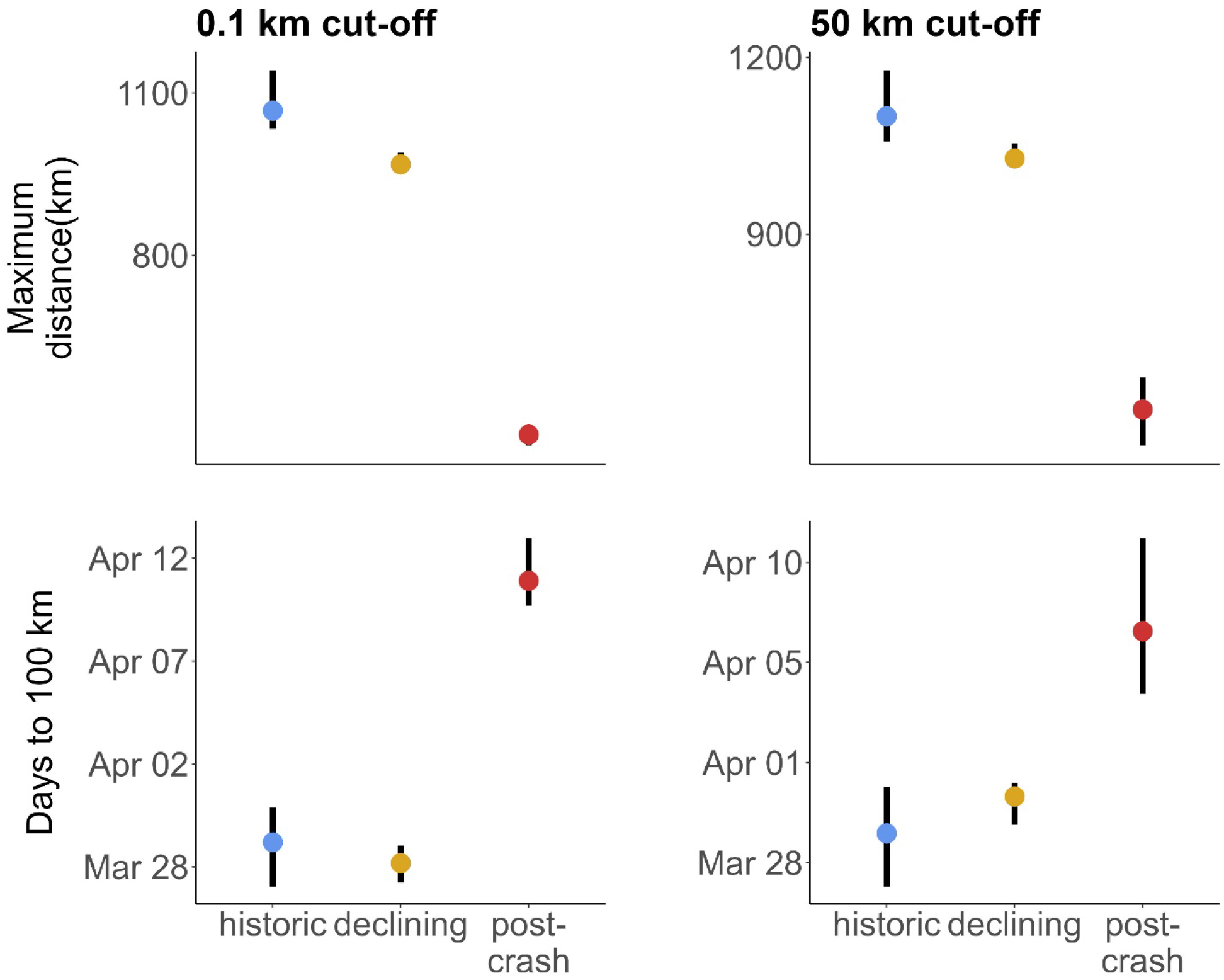
Qualitative patterns of seasonal range expansion are insensitive to choice of threshold distance for excluding observations near the overwintering sites. We repeat our in the quantile choice. We repeat our analyses and replicate Fig. 1F and 2C for 50 km or 0.1 km threshold distances (as compared to 30 km in the main text).

**Supplemental Figure S4:**
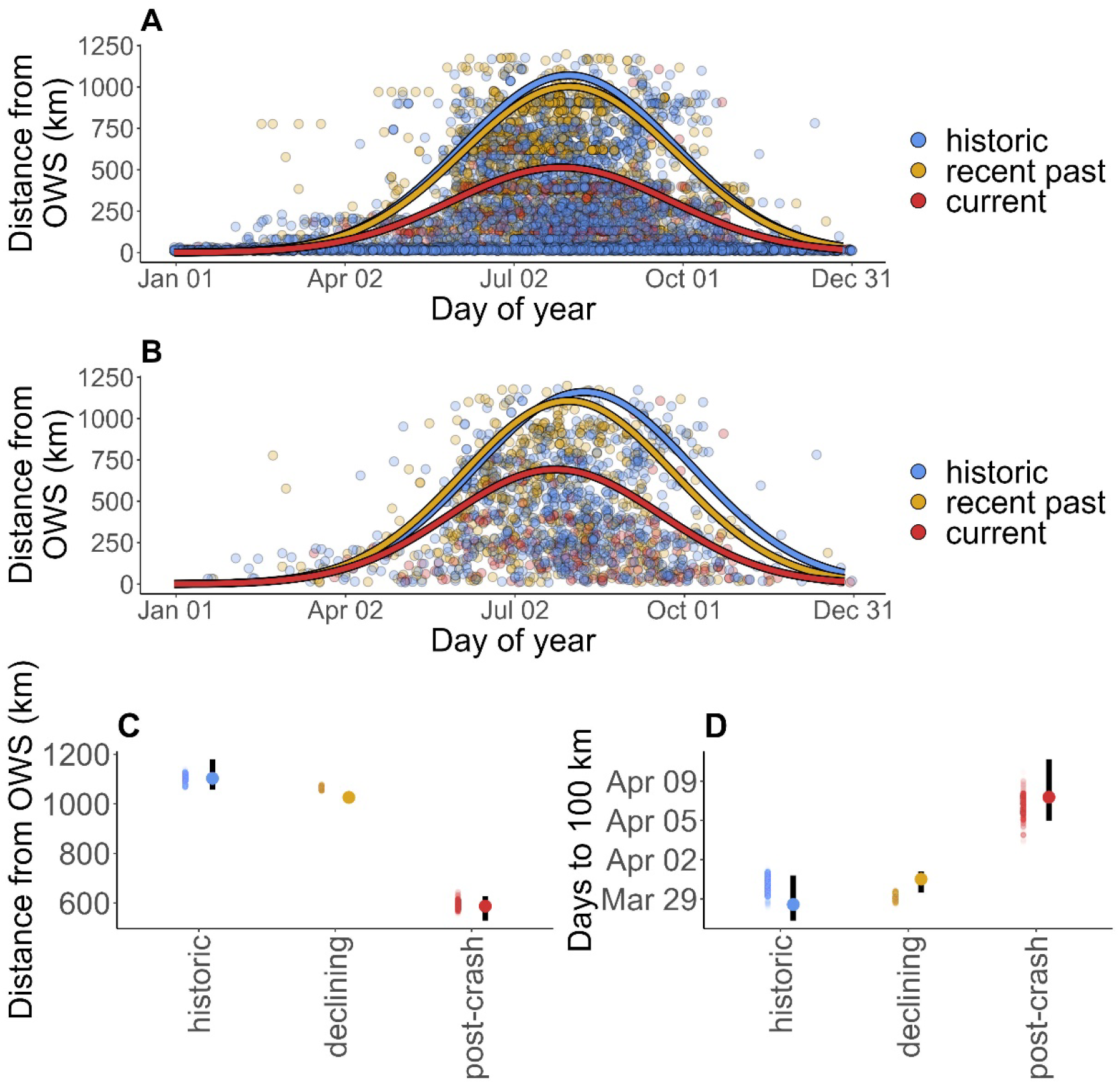
(A) Data showed apparent horizontal “streaks” of multiple data points at the same distance from overwintering sites, potentially representing individual locations with frequent sampling. To ensure dynamics of these locations were not biasing our results, we simulated data by randomly sampling only one observation per era per kilometer of distance from the overwintering site (i.e., “data thinning”). (B) Example of this simulated data. C-D: range and phenology metrics calculated from 5000 simulated datasets (cloud of points, left) qualitatively reflected the estimates from the full data set (point and bars, right, same as Fig. 1F-G).

